# Continuous hypermutation and evolution of luciferase variants

**DOI:** 10.1101/2025.08.11.669707

**Authors:** Tanya Hadjian, Ria A. Deshpande, Zachary R. Torrey, Amelia W. Hammond, Rosana S. Molina, Kevin Ng, Chang C. Liu, Jennifer A. Prescher

**Affiliations:** Department of Chemistry, University of California, Irvine, CA, USA; Program in Mathematical Computational and Systems Biology, University of California, Irvine, CA, USA; Department of Biomedical Engineering, University of California, Irvine, CA, USA; Center for Synthetic Biology, University of California, Irvine, CA, USA; Department of Pharmaceutical Sciences, University of California, Irvine, CA, USA; Department of Molecular Biology & Biochemistry, University of California, Irvine, CA, USA

## Abstract

Several luciferases have been developed for imaging and biosensing, and the collection continues to grow as new applications are pursued. The current workflow for luciferase optimization, while successful, remains laborious and inefficient. Mutant libraries are generated in vitro and screened, “winning” mutants are picked by hand, and the isolated sequences are subjected to additional rounds of mutagenesis and screening. Here, we present a streamlined platform for luciferase engineering that removes the need for manual library generation during each cycle. We purposed an orthogonal DNA replication (OrthoRep) system for continuous hypermutation of a well-known luciferase (GeNL). Short cycles of culturing and screening were sufficient to evolve the enzyme, with no repetitive manual library generation necessary. New GeNL variants were identified that exhibit improved light outputs with a non-cognate and inexpensive luciferin. We further characterized the novel luciferases in cell models. Collectively this work establishes OrthoRep and continuous hypermutation as a viable method to engineer luciferases, and sets the stage for more rapid development of bioluminescent reporters.

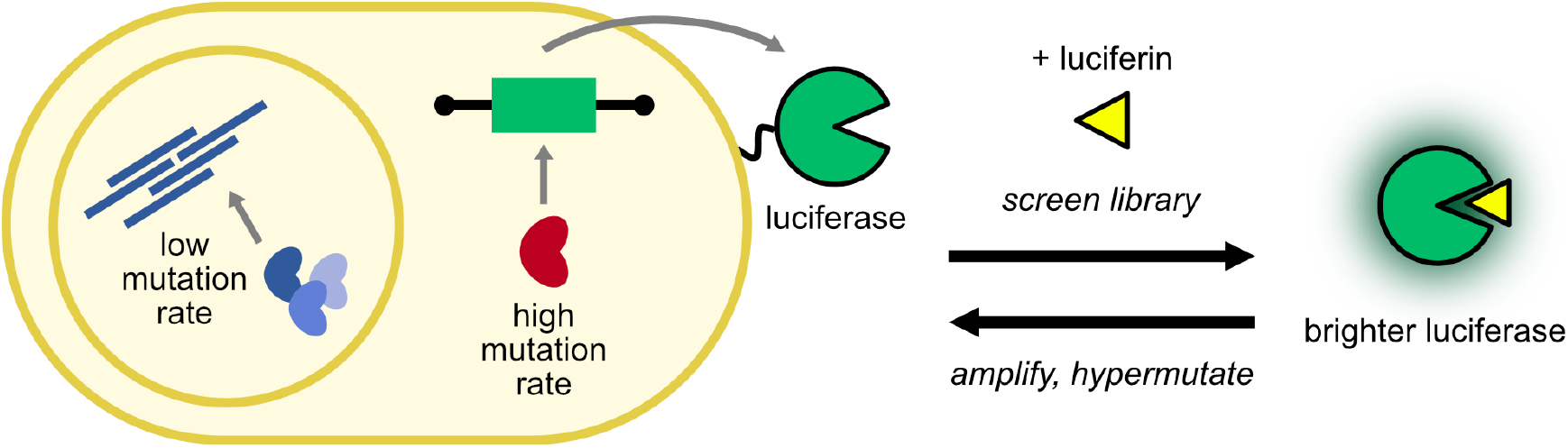

## Introduction

Bioluminescence is used extensively for sensitive imaging in vitro and in vivo.^1–4^ At the heart of this technology are robust luciferase enzymes that emit photons with small molecule luciferins. Bioluminescence does not require external excitation light, making it advantageous for probing biological systems over time and in complex environments. The breadth of applications has expanded dramatically in recent years, thanks to the development of designer luciferases and luciferins.^5,6^ Dozens of unique bioluminescent tools are now available, with many exhibiting unique emission profiles, stabilities, and substrate preferences (Figure 1A).^7–10^ This number will also likely continue to climb, as the demand for excitation-free imaging increases and bioluminescent cameras become more accessible.

**Figure 1.**
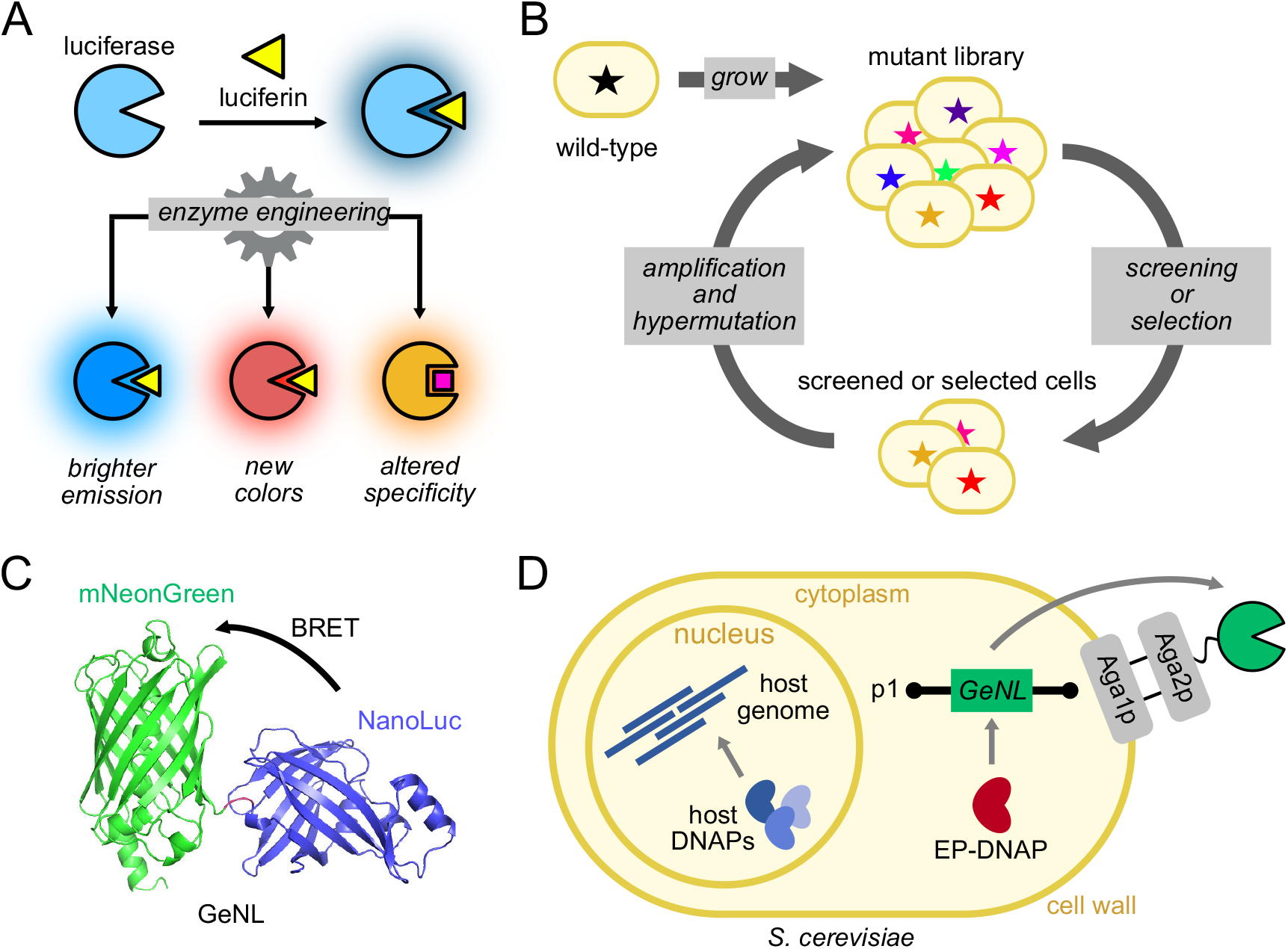
Streamlined evolution of luciferases by OrthoRep. (A) General bioluminescence reaction. A luciferase enzyme oxidizes a small-molecule luciferin to produce light. Luciferase engineering can provide enzymes with novel features. (B) General scheme for continuous evolution. Evolution cycles can be carried out quickly in cells, since autonomously generated mutant libraries are repeatedly screened or continuously selected for desired function. (C) Luciferase used in this study. GeNL exhibits bioluminescence resonance energy transfer (BRET) with NanoLuc as the donor and mNeonGreen as the acceptor. (D) Engineered OrthoRep AHEAD yeast strains allow for surface expression of GeNL.

Identifying new luciferases to support the bioluminescence technology ecosystem has historically been quite labor-intensive.^11–14^ The process typically involves generating large mutant libraries via error-prone PCR or other manual cloning techniques. The libraries are then screened using various luciferins. Light emission is very weak, precluding standard selections or rapid sorts with common instruments (e.g., flow cytometry). Instead, screens are conducted via manual assessment of individual colonies on agar plates. Bacteria are often used as large amounts of each luciferase must be produced for signal detection. Screening of each hit typically involves spraying or layering concentrated solutions of luciferin substrates on agar plates, which can be expensive in large volumes.^13-17^ “Hits” identified each cycle are isolated and sequenced. This process then repeats with iterative cycles of manual library generation, transformation, and screening.

We aimed to accelerate the process of luciferase discovery using systems for continuous evolution (Figure 1B).^18^ This strategy relies on targeted hypermutation systems in engineered cells. Continuous hypermutation of a gene of interest can provide similar or greater coverage as conventional error-prone methods, but with minimal cloning and cell manipulations required, as the mutagenesis automatically occurs in vivo.^19^ We reasoned that this approach could ease one of the bottlenecks in luciferase discovery by removing the need to retrieve, characterize, and re-diversify top variants each round. We were particularly drawn to OrthoRep, a platform based on an orthogonal error-prone DNA replication system that rapidly and durably mutates target genes in yeast.^20^ OrthoRep features an error-prone polymerase (EP-DNAP) that specifically replicates a multicopy plasmid (p1) encoding a gene of interest. Importantly, EP-DNAP replicates p1 through a protein-primed replication process in the cytoplasm, providing mechanistic and physical separation from host genome replication so that the host genome does not experience elevated mutation rates. Optimized variants of the EP-DNAP mutate p1 at rates ∼10^5^-10^6^ times higher than the mutation rate of the host genome, resulting in ∼1 mutation in a typical 1 kb gene every ∼10 generations.^21^ OrthoRep has been successfully used to evolve several protein targets, including synthases^22^ and antibodies.^23–25^ Notably, the platform is compatible with surface-displayed target proteins.^23,24^ This feature is advantageous for luciferase engineering, as many luciferins have suboptimal cell permeabilities.

Here we report the application of OrthoRep to luciferase evolution. We established a yeast strain for continuous hypermutation of a luciferase-fluorescent protein fusion (GeNL). The bioluminescent reporter was further fused to Aga2 for surface display. Using OrthoRep, GeNL was evolved for improved processing of a cost-effective luciferin (coelenterazine). New variants were identified, comprising mutations that have not been seen in previous evolution campaigns. Further biochemical analyses and cell assays were performed to validate the GeNL mutants. Collectively, this work showcases a new platform for targeted luciferase evolution and discovery.

## Results

### Design of a luciferase evolution system using the yeast OrthoRep system

As a starting point, we selected the green enhanced Nano-lantern (GeNL) for continuous hypermutation. GeNL comprises a fluorescent protein, mNeonGreen, fused to NanoLuc luciferase (Figure 1C).^26^ GeNL is one of the brightest bioluminescent probes reported to date and has been leveraged for length scales ranging from subcellular to whole animal imaging.^27,28^ When incubated with luciferins, the normal blue emission of NanoLuc shifts to green via bioluminescence resonance energy transfer (BRET). The optimized substrate for GeNL is furimazine (Fz), a synthetic analog of coelenterazine (CTZ).^29^ Fz generates more robust signal with GeNL and other NanoLuc-based luciferases compared to CTZ.^29,30^ However, CTZ, the native substrate for many marine luciferases (including the one from *Oplophorus gracilirostris* from which NanoLuc is derived), is less expensive than Fz. More efficient CTZ-utilizing luciferases are thus desirable to achieve bright photon outputs at minimal cost. Excitingly, structure-guided mutagenesis recently revealed an allosteric NanoLuc mutant (NanoLuc^CTZ^) with ten-fold improved processing of CTZ.^30^ We aimed to discover novel GeNL variants capable of robust bioluminescence with CTZ to determine the capabilities of the OrthoRep platform for luciferase evolution. We first needed to establish the starting yeast strain for evolution (i.e., GeNL encoded in the p1 plasmid, Figure S1). Toward this end, we selected an OrthoRep strain encoding a highly error-prone orthogonal DNA polymerase (EP-DNAP), BadBoy2, for replication of the p1 plasmid during cell proliferation. BadBoy2 has an error rate of 1.4 × 10^−4^ substitutions per base.^21^ Additionally, we decided to encode GeNL as a surface-displayed protein, similar to previous work with antibody targets encoded on p1 in AHEAD strains^23^ (Figure 1D). Surface display is desirable in cases where substrate permeability is limiting, as can be the case with some luciferins. AHEAD yeast strains feature an agglutinin subunit (Aga2) fused to the protein target, all encoded on p1. The anchor subunit, Aga1, is encoded in the genome under the control of a *GAL1* promoter (*pGAL1*). Galactose-induced Aga1 expression ultimately enables display of the target-Aga2 fusion on yeast cell surfaces.

We thus designed a linear p1 plasmid encoding GeNL fused to the N-terminus of Aga2, using a strong p1-specific constitutive promoter (p10B2).^31^ Galactose induction would thus result in surface display of GeNL, enabling easy access of the CTZ substrate. The desired plasmid was constructed via integration of a GeNL donor cassette into p1. Successful integration was confirmed upon transformation (Figure S2). We further verified GeNL expression upon galactose treatment. Induced cultures incubated with CTZ showed a two-fold increase in luminescence (Figure S3). Light emission from the uninduced cultures was attributed to intracellular GeNL.

### Screening luciferase libraries in OrthoRep

After the GeNL target sequence was successfully integrated into OrthoRep yeast, we began the repetitive cycle of hypermutation and screening (Figure S4).^32^ An initial culture was grown from a starting population (P0) and passaged serially to allow for mutations in p1 to accumulate. Cultures were then plated and colonies were picked to yield passage 1 (P1). A total of 94 colonies were picked in passage 1 and grown in deep-well plates. The resulting populations were assigned an arbitrary number from 1-94 and then induced for surface display. CTZ was administered and luminescence was recorded. The brightest populations for each passage were identified and plated to single colonies, and ∼10 colonies were used to inoculate fresh cultures. The process was repeated for seven total passages (<2 months).

The imaging results from P1 showed large variations in emission intensity across the individual populations, suggesting that different GeNL variants were indeed being generated through hypermutation. Mutant population 72 (referred to as P1-72) was particularly noteworthy for its brightness. This culture produced roughly 5-fold brighter signal than the next top five populations (Figure 2A). The top populations in all subsequent cycles of evolution originated from P1-72, as we diagram in a map of lineages of the top performing populations (Figure 2B). Once the screening cycles were complete, a secondary luminescence analysis was performed for the top-performing mutant populations from each plate. Each of the selected populations exhibited 3- 10 fold improvements in photon output with CTZ compared to the initial P0 culture (Figure 2C). We further performed a separate evolution experiment with OrthoRep and the same target GeNL sequence. Improvements in light output with CTZ were also observed with increasing passage number (Figure S8A), highlighting the overall reproducibility of the approach.

**Figure 2.**
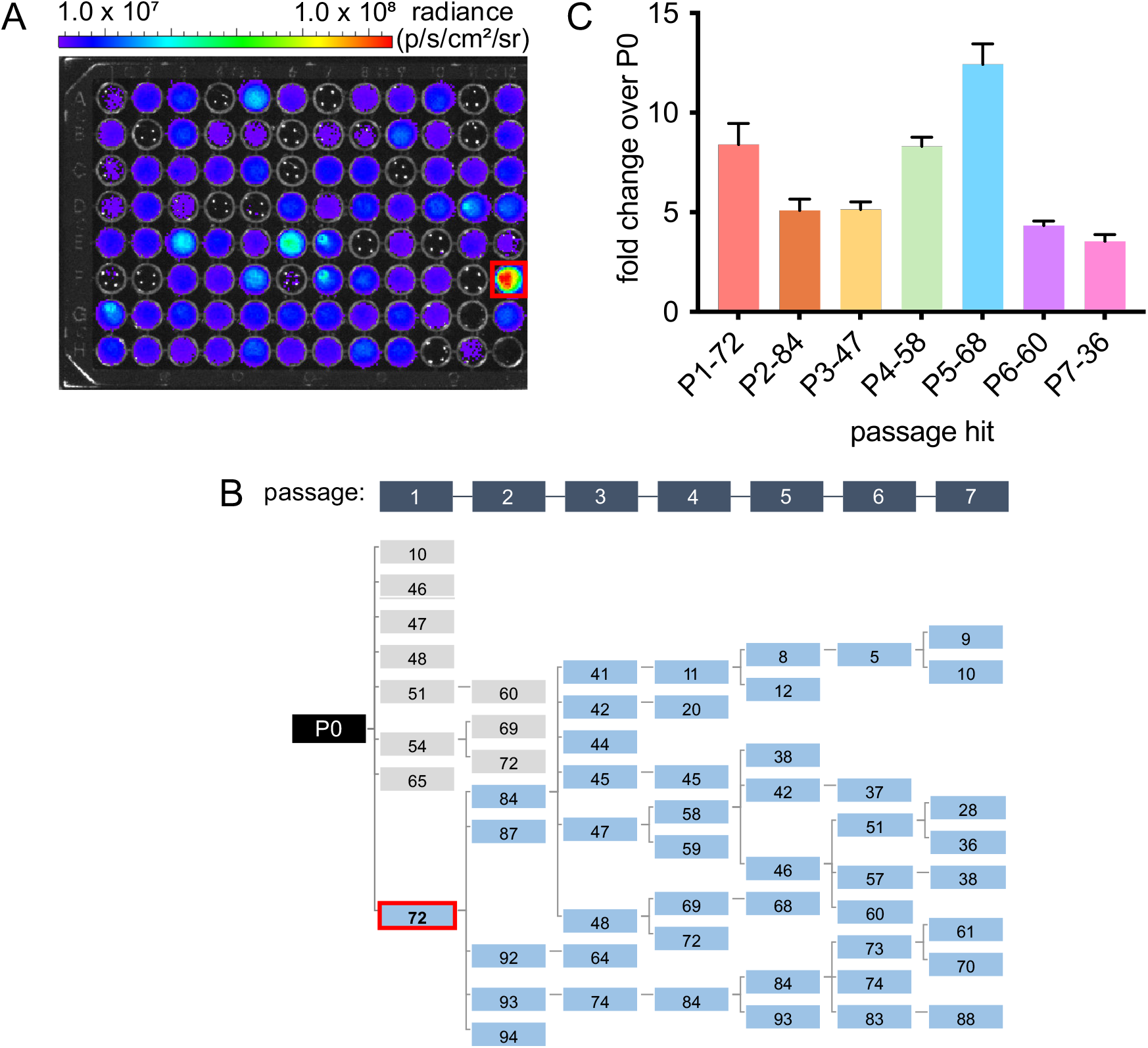
Screening revealed bright GeNL populations. (A) Representative image of passage 1 populations screened and imaged with CTZ in a black 96-well plate. (B) Phylogeny tree showing the origin of the top 8 populations from each passage. (C) Photon outputs for the top 8 populations compared to GeNL-expressing yeast. Data are plotted as fold change values over passage 0. Error bars represent the standard error of the mean for *n* = 3 technical replicates.

### Biochemical analyses of top variant shows improvement over GeNL

We identified the mutations present in each population selected from the OrthoRep evolution campaign (Supplementary Table S1). Three coding mutations were found in the brightest mutant from each passage (Figure 3A). One coding mutation (D147Y) was found in mNeonGreen, while two mutations (L243I, H312Y) were present in NanoLuc (Figure 4B). We also identified a fourth mutation in our second evolution campaign (I283T). This mutation was also present in the luciferase region.

**Figure 3.**
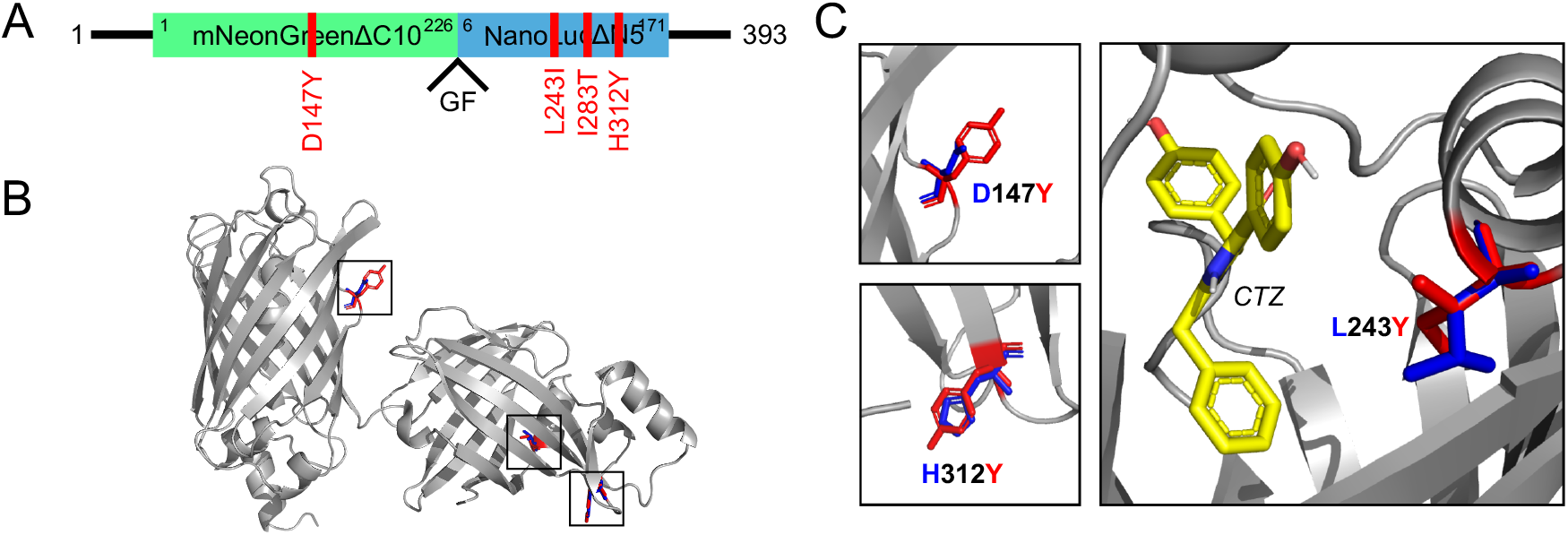
Four mutations from OrthoRep-driven evolution experiments were prioritized for characterization. (A) Mutations located in GeNL on p1. (B) Structure of GeNL with wild-type residues (blue) overlaid with mutations found during evolution (red). GeNL model was generated by AlphaFold2. (C) Close-up views of mutated residues and proximity to bound CTZ.

**Figure 4.**
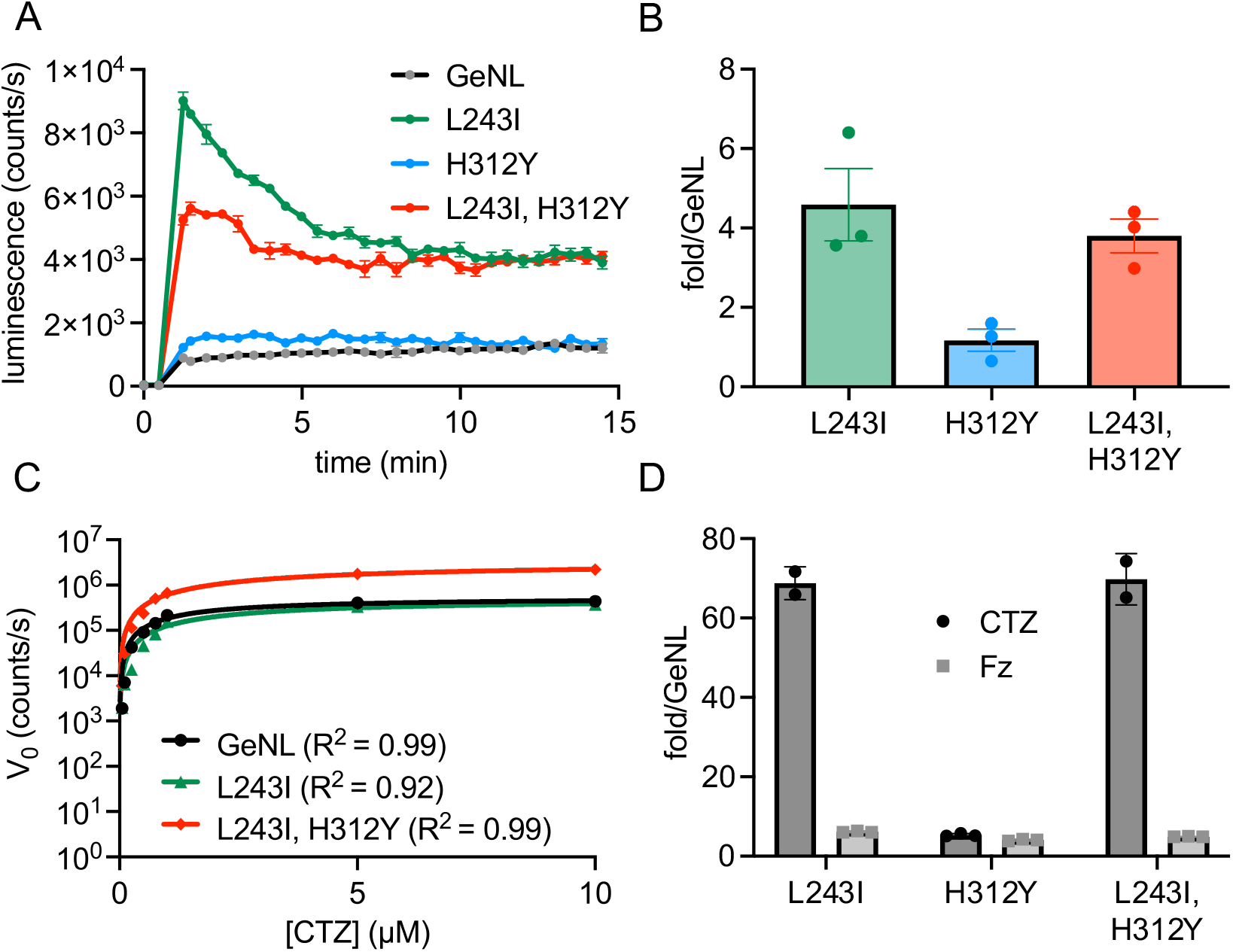
Characterization of GeNL mutants. (A) Cloned yeast variants were treated with CTZ (at t = 2 min) and imaged over time. Error bars represent the standard error of the mean for *n* = 3 replicate measurements, and the data are representative of *n* = 3 biological replicates. Luminescence was normalized to mNeonGreen fluorescence to account for differences in expression. (B) Fold change in luminescence for cloned yeast variants 15 min post-CTZ treatment. Error bars represent the standard error of the mean for *n* = 3 biological replicates. (C) Determination of kinetic constants for purified mutants and CTZ. The calculated *K*_m_ values are apparent and were determined via measurements of initial rates of light emission over a range of CTZ concentrations (0–10 µM). Error bars represent the standard error of the mean for *n* = 3 replicates. (D) Fold change in luminescence for purified mutants treated with CTZ (20 µM) or Fz (10 µM). Error bars represent the standard error of the mean for *n* = 3 measurements.

We focused on two of the four mutations (L243I, H312Y) in follow up biochemical assays. Residues 243 and 312 are both localized to luciferase and closest to the putative luciferin binding site (Figure 3C and S5). Docking studies by Altamash *et al*. suggested that L243 (equivalent to residue L18 in NanoLuc) mediates hydrophobic interactions with the luciferin substrate (Fz).^33^ The proximal location of L18 within the binding pocket was further corroborated by crystal structure analyses of NanoLuc (containing a bound luciferin analog).^30^ The functional role of the L18 residue was not experimentally verified in either case, though. We further analyzed the L243I mutant using PyMOL and AutoDock Vina (Figure 3C). The size of the binding pocket decreased compared to NanoLuc, suggesting a potential mechanism for improved CTZ utilization and overall brightness.^33^

We next examined the functional impact of L243I and H312Y mutations, individually and in combination, on GeNL in yeast. Variants were encoded on a non-mutating nuclear CEN/ARS plasmid and their expression was driven by a weak promoter, pREV1. To eliminate any potential variability in enzyme display, we expressed the luciferases cytosolically. CTZ was administered to the cultures and light emission was recorded. The mutants exhibited ∼5-fold more robust emission compared to GeNL (Figures 4A-B). We also measured the luminescence emission spectra of each mutant. The mutations did not significantly alter energy transfer between NanoLuc and mNeonGreen. As shown in Figure S6, the bioluminescence emission spectra of all mutants matched that of GeNL with CTZ.

We further examined the luciferases in vitro. Both the mutants and GeNL were purified using standard bacterial expression systems (Figure S7). Kinetic assays were performed with varying amounts of CTZ. The apparent *K*_m_ values for the mutants and GeNL were similar. However, the mutants exhibited higher V_max_ values (Figure 4C, Supplementary Table 2). No significant differences in brightness with Fz were noted between the mutants and GeNL (Figure 4D).

The top performing mutant was tested in mammalian cells. HEK293 cells were engineered to express the L243I mutant or GeNL itself. The cultured cells were then incubated with CTZ and imaged over time. As shown in Figure 5A-B, the mutant provided improved photon outputs at earlier time points. Sustained emission on par with GeNL was observed at later time points.

**Figure 5.**
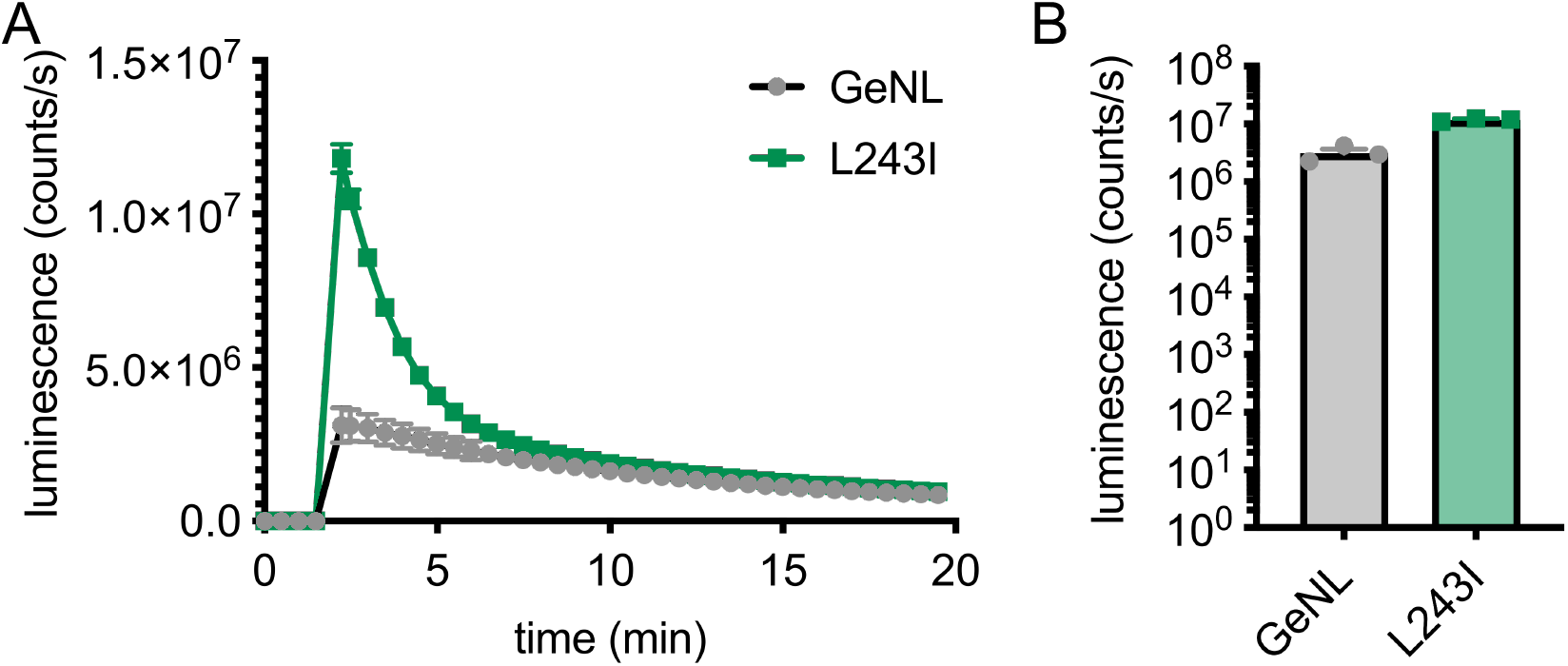
Cell imaging with L243I mutant and CTZ. (A) GeNL or mutant-expressing HEK cells treated with CTZ (2 µM, administered at t = 2.5 min) and imaged over time. Luminescence values were normalized to mNeonGreen expression. (B) Maximum luminescence values for cells treated with CTZ as in (A). Error bars represent the standard error of the mean for *n* = 3 replicate measurements.

## Conclusions

Here, we used OrthoRep as a rapid evolution platform to evolve a brighter GeNL variant with coelenterazine. OrthoRep enables gene hypermutation in yeast, providing an easy-to-use platform for evolving luciferases. As an initial evaluation of the platform, we generated a strain of yeast capable of expressing and mutating GeNL. The luciferase was evolved over seven cycles of screening, with mutants from each cycle originating from the one before it. After a few weeks, the brightest hits were isolated and characterized in multiple expression systems. The winning mutants exhibited improved utilization of a model luciferin and could be used for imaging in model cell lines.

We expect OrthoRep to be useful for evolving additional luciferases. OrthoRep eliminates the need for multiple rounds of cloning and transformation, shortening the time to discover new enzyme variants. Yeast surface display is also beneficial in cases where luciferin permeability is limiting, which can be impeding in other screening platforms with bacteria. New versions of AHEAD strains will further streamline the process.^24^ These new strains feature a β-estradiol induction system, which enables surface display of the protein of interest in as little as one hour, in contrast to the standard two-day galactose induction system used here. The shorter induction times will vastly reduce the time required per evolutionary cycle.

It should be noted that the luciferase evolution platform described here was likely limited by population size and throughput, as only ∼96 clones and populations were screened per cycle, which is much less than the expected mutant diversity that OrthoRep was generating per cycle. A high-throughput screening approach, such as cell sorting or growth-based selection, is needed to exploit the continuous diversification speed and scale of OrthoRep. However, luciferase emission is inherently dim, capturing signals on a single-cell level during sorting is challenging, and no commercially available instruments exist for luminescence sorting. Fluorescence-activated cell sorting (FACS) could be used but requires activity that can be linked to a fluorescent reporter, which cannot be directly applied here. Future work will be directed at addressing these shortcomings in screening throughput, which will continue to relieve the historic bottlenecks in luciferase evolution and discovery that this work has begun to remove.

## Acknowledgements

This work was supported by the Paul G. Allen Frontiers Group (to J.A.P.) This project was further supported by the U.S. National Institutes of Health (NIH) R35GM136297 (to C.C.L.). Flow cytometry analysis was performed at the Institute for Immunology Flow Cytometry Core (UCI). We also thank members of the Prescher and Liu labs for helpful discussions, including Dr. Alon Wellner for sharing materials and advice.

## Notes

### Competing Interest Statement

The authors have declared no competing interest.

