## Supplementary Materials for "Continuous hypermutation and evolution of luciferase variants"

### TABLE OF CONTENTS

| | |
|---|---|
| Figures S1-S8 | S2-S9 |
| Materials and Methods | S10 |
| Supplementary Tables | S14-S18 |
| Supplementary Notes | S19-S20 |
| References | S21 |

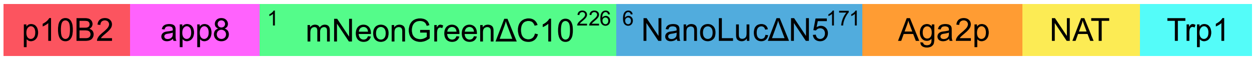


**Figure S1.** Schematic of linear integrated p1 plasmid containing GeNL (mNeonGreenΔC10-NanoLucΔN5 fusion) and other notable components.

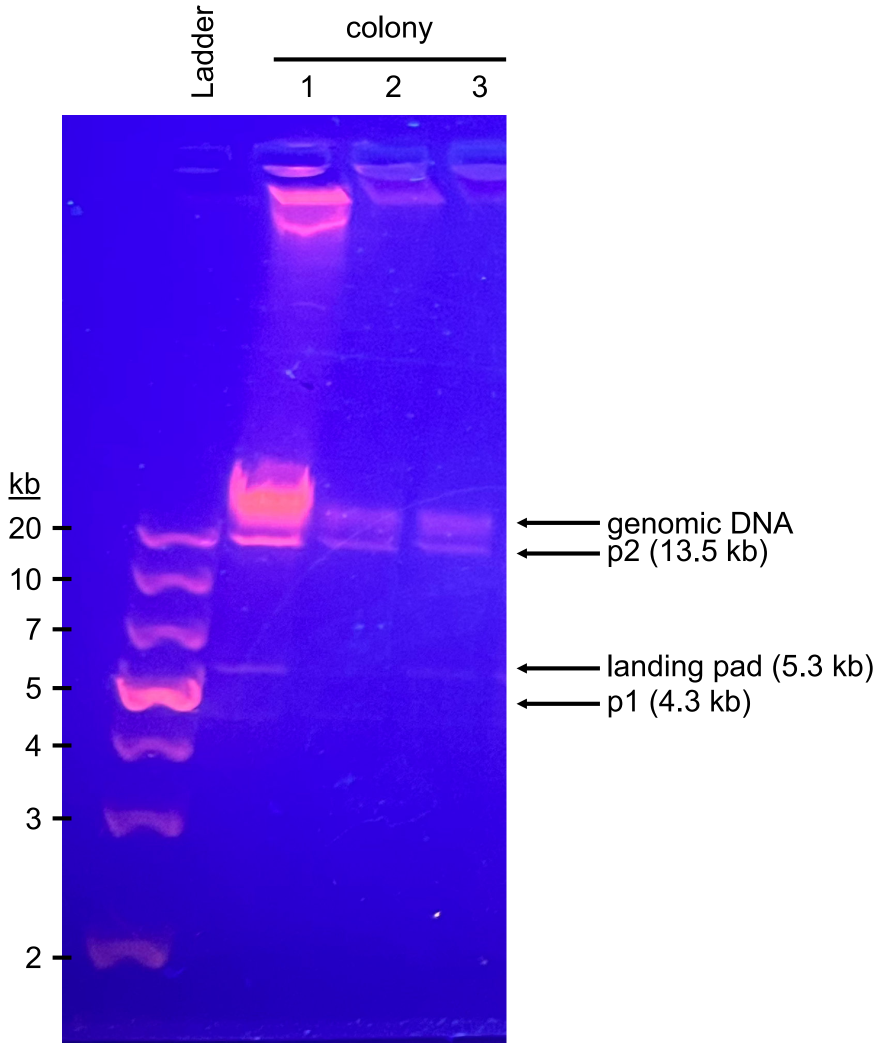


**Figure S2. Confirmation of p1 integration.** DNA was extracted from three randomly selected colonies (following integration of the GeNL gene) and analyzed via gel electrophoresis.

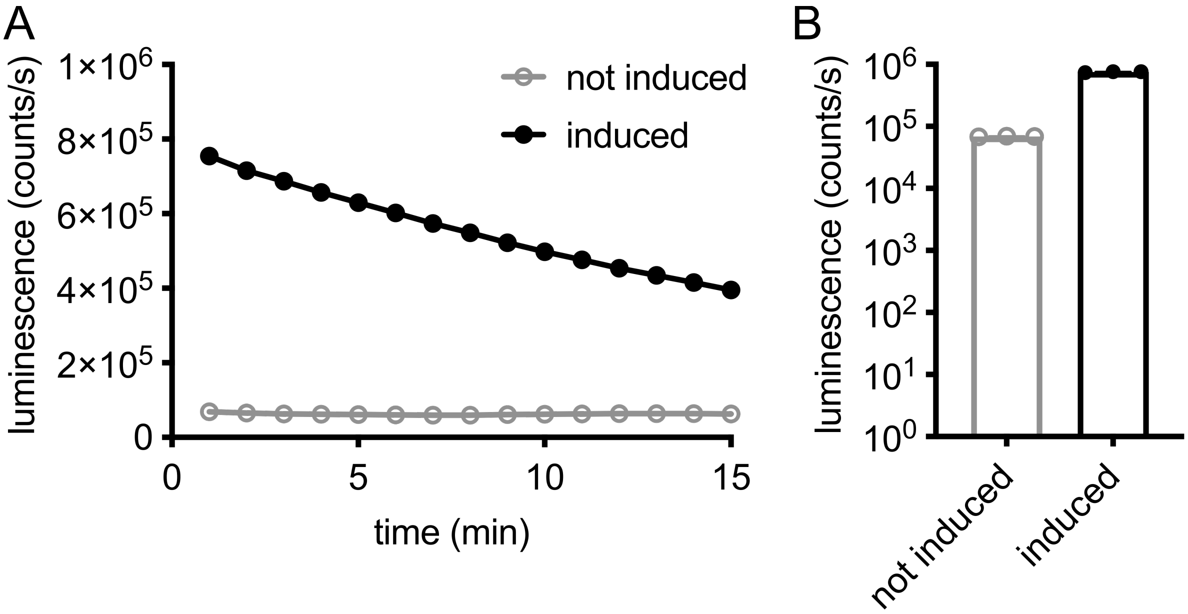


**Figure S3. Surface-display induction increases bioluminescence emission.** (A) Normalized luminescence emission of induced and non-induced starting culture of GeNL-integrated yeast. Cultures were incubated with CTZ (5 μM final concentration) and monitored for 15 min. (B) Maximum normalized luminescence emission for each condition. In (A)-(B), error bars represent the standard error of the mean for *n* = 3 replicate measurements.

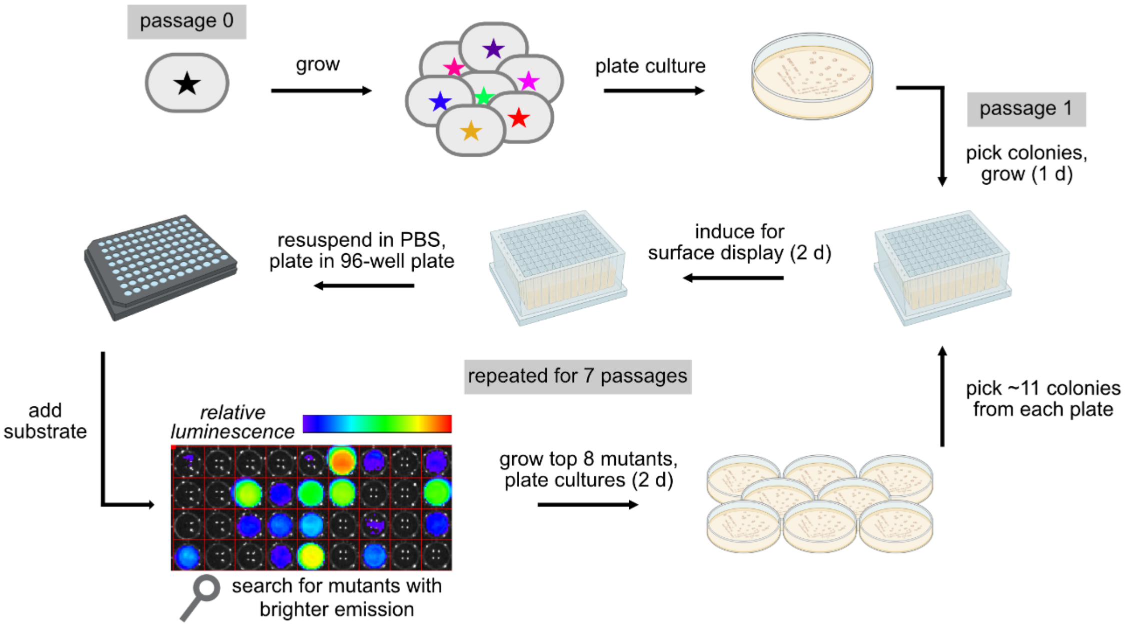


**Figure S4. General workflow for OrthoRep evolution of GeNL.** The GeNL gene was incorporated into p1-expressing yeast, and the starting culture (“passage 0”) was plated. Colonies were randomly picked to fill a 96-well plate. Colonies were grown to saturation, then resuspended in media containing 2% galactose to induce surface display. After 2 days of shaking at room temperature, cultures were resuspended in PBS and plated in a black 96-well plate. Substrate (CTZ) was administered to each well and images were acquired. The screening cycle was repeated for 7 total rounds. Figure prepared using BioRender.

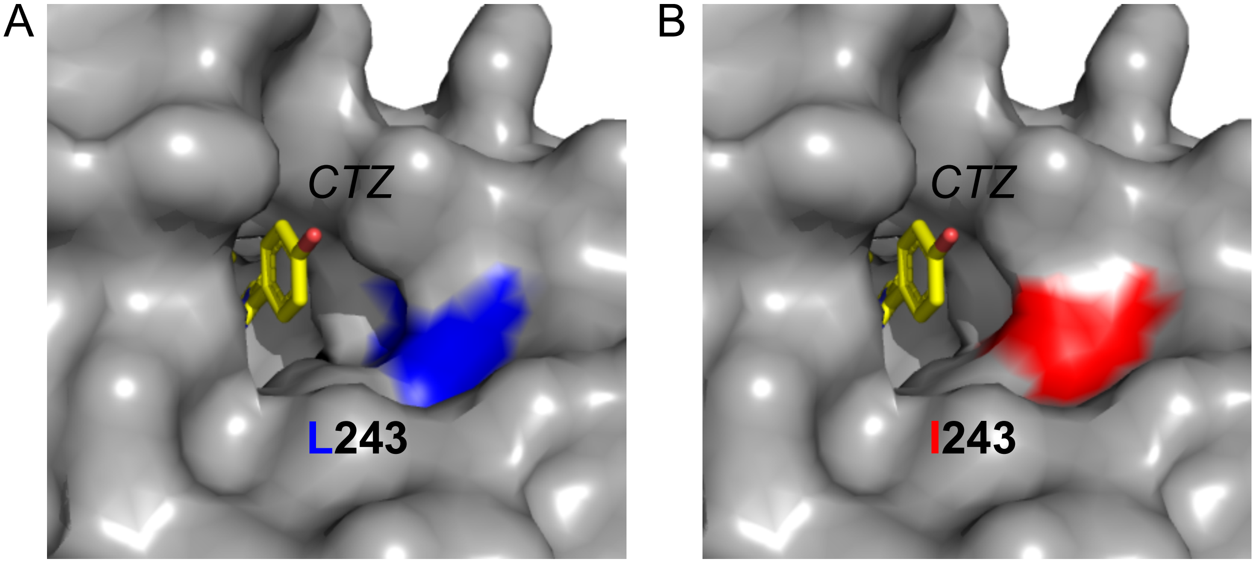


**Figure S5. Molecular docking of CTZ with NanoLuc and L243I mutant active site.** CTZ was docked into NanoLuc (PDB: 5B0U) using AutoDock Vina. (A) Residue equivalent to L243 in GeNL (blue) shown in NanoLuc. (B) Residue equivalent to I243 in GeNL (red) shown in the mutated NanoLuc.

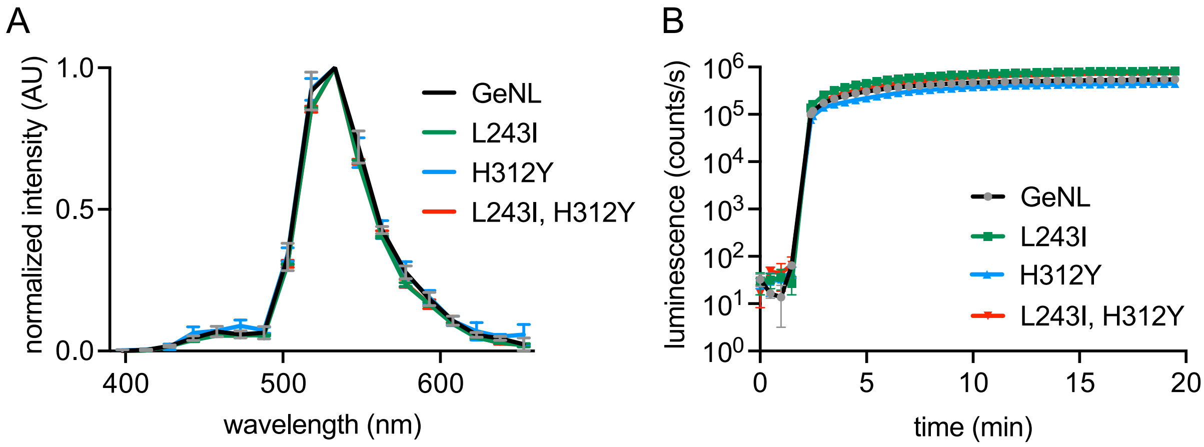


**Figure S6. Bioluminescence emission of mutants expressed in yeast.** (A) Mutants expressed in CEN/ARS yeast strains were treated with CTZ and imaged. (B) Yeast cultures were treated with Fz and imaged over time. In (A)-(B), one representative experiment from 3 biological replicates is shown. Error bars represent the standard error of the mean from $n$ = 3 measurements.

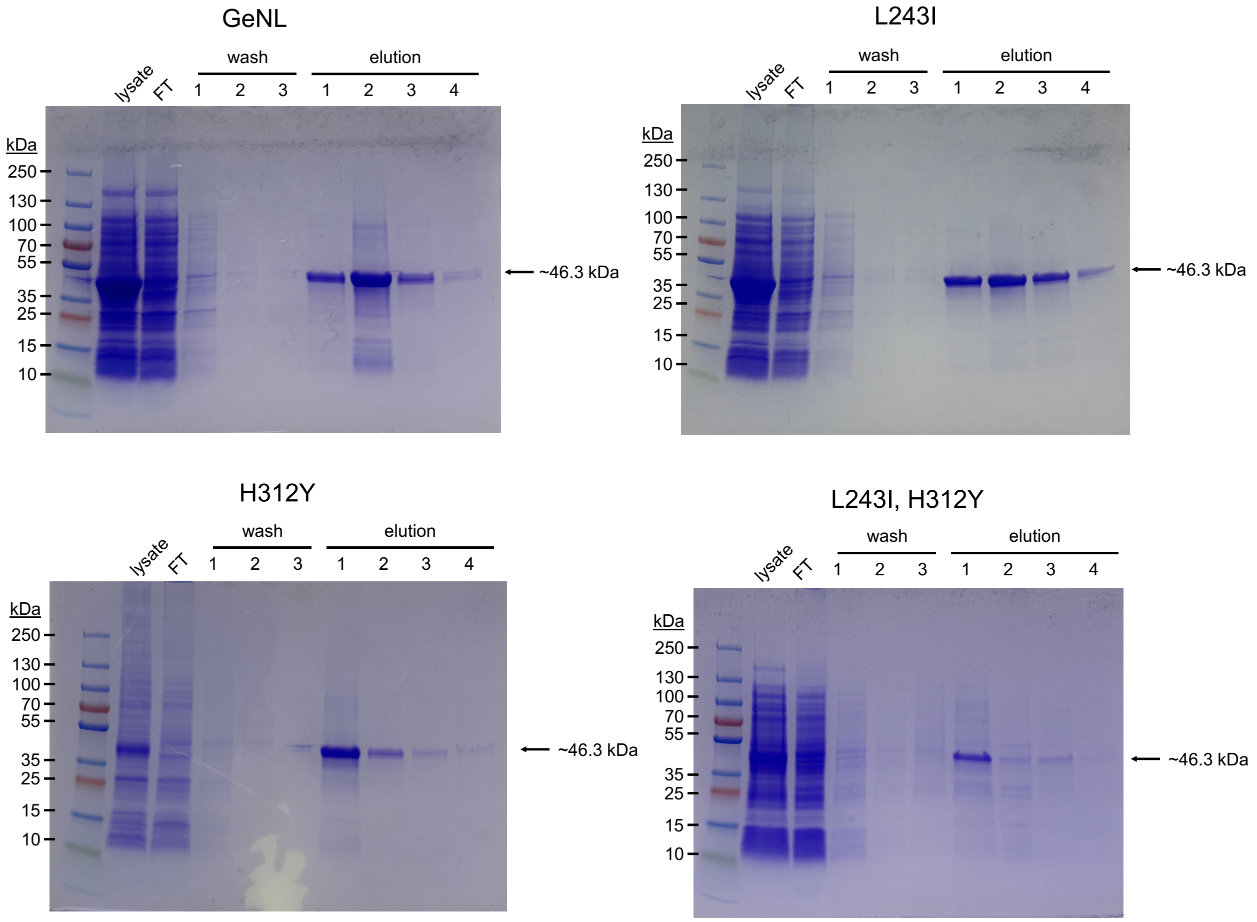


**Figure S7. SDS-PAGE confirming purity of purified recombinant proteins.** Proteins were expressed in *E. coli* BL21 cells and purified by nickel-affinity chromatography. SDS-PAGE gels were stained with Coomassie Brilliant Blue R-250.

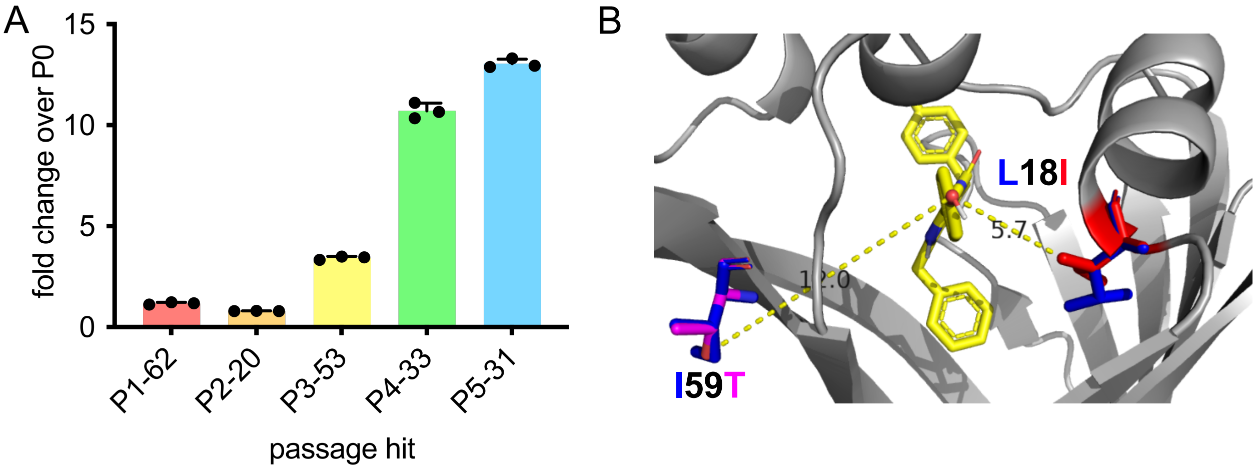


**Figure S8. A second OrthoRep evoultion of GeNL.** (A) GeNL-expressing yeast were subjected to similar growth and screening cycles as in Figure 2. Relative luminescence values for the top populations in each passage are shown. (B) L18I and I59T mutations shown in NanoLuc, along with docked CTZ.

### Materials and Methods

#### Materials

Q5 DNA Polymerase and restriction enzymes were purchased from New England Labs. dNTPs were purchased from Thermo Fischer Scientific. Luria-Bertani medium (LB) was purchased from Genesee Scientific. NanoGlo luciferase substrate (furimazine, Fz) was purchased from Promega. Coelenterazine (CTZ) was purchased from GoldBio. All plasmid and primer stocks were stored at –20 °C.

#### General cloning methods

Polymerase-chain reaction (PCR) was performed to amplify and isolate genes of interest. The yeast-codon optimized GeNL was synthesized as a gBlock (Integrated DNA Technologies, IDT). Primers were purchased from Integrated DNA Technologies (IDT). Plasmids and PCR products were sequenced by Genewiz from Azenta Life Sciences and analyzed using Benchling. Plasmids were assembled using Gibson assembly[1] or Golden Gate[2] cloning, where specified. PCR reactions, Gibson Assemblies, and Golden Gate Assemblies were performed in a Bio-Rad C1000 Touch Thermal Cycler.

#### Plasmid construction

All plasmids, yeast strains, and primers are described in Supplementary Tables S3, S4, and S5, respectively.

An integration plasmid to insert GeNL into p1 was constructed (RM-Ec033). RM-Ec033 was made by replacing the nanobody coding sequence in plasmid pAW463 (equivalent to plasmid pAW240 (Addgene #170791)[3] but with a different nanobody sequence) with a codon optimized GeNL gene using Gibson assembly. The backbone fragment for Gibson assembly was generated from RM-Ec033 via restriction digestion with *Kpn*I and *Sal*I instead of PCR in order to preserve a functional poly-A sequence that is easily shortened during PCR.

For expression in bacteria, GeNL was cloned into a pCold backbone. The pCold vector was digested with *Nde*I and *Eco*RI. The GeNL insert was amplified from a pcDNA-GeNL plasmid (Addgene#85200; GeNL/pcDNA3) with primers TAH06_Fwd and TAH07_Rev. The following thermal cycling conditions were used to amplify the GeNL insert: 20 cycles of denaturation (95 ˚C, 30 s), annealing (65 ˚C, 30 s) and extension (72 ˚C, 90 s). Site-directed mutagenesis (SDM) with partially overlapping primers was performed to obtain the mutated variants using pTAH003 as a template. The following thermal cycling conditions were used for SDM: 20 cycles of denaturation (95 ˚C, 30 s), annealing (63 ˚C, 30 s) and extension (72 ˚C, 6 min).

To obtain plasmids for expression of the variants in yeast, variants were cloned into a CEN/ARS backbone. The plasmid pAW729 was digested with *Bgl*II and *Xho*I to obtain the CEN/ARS vector.[4] The necessary gene inserts for GeNL and the variants were built using Golden Gate assembly. Segments of GeNL were amplified from plasmid RM-Ec033 using PCR with appropriate primers to introduce mutations. The following thermal cycling conditions were used to amplify the GeNL fragments: 20 cycles of denaturation (95 ˚C, 30 s), annealing (63 ˚C, 30 s) and extension (72 ˚C, 90 s). GeNL and the variants were assembled using Golden Gate assembly with custom primers into the entry vector (pYTK001) from the yeast toolkit.[5] Following entry vector assembly, the full inserts were amplified using primers TAH029_Fwd and TAH030_Rev. Gibson assembly was used for final construction of the CEN/ARS plasmids.

For mammalian cell expression of the L243I variant, a pcDNA-GeNL plasmid (Addgene, plasmid #85200) was digested with *Bam*HI and *Eco*RI to obtain the pcDNA vector. The GeNL-L243I insert was amplified from pTAH004 with primers ZY28 and ZY29. The following thermal cycling conditions were used to amplify the GeNL insert: 20 cycles of denaturation (95 ˚C, 30 s), annealing (68 ˚C, 30 s) and extension (72 ˚C, 75 s). Gibson assembly was performed to obtain plasmid pTAH061.

**Yeast transformation**

Plasmids were transformed into yeast strains to incorporate the target genes into p1 using the high-efficiency Gietz yeast transformation method. [6] In brief, starter cultures of yeast were grown to saturation in a selective SC media minus the appropriate amino acids (SC-HLUMC for yAW723, SC-HUMC for yAW680). 330 µL of saturated culture was used to inoculate 10 mL of YPD medium, which was grown to mid-log phase at 30 °C. The cells were pelleted by centrifugation (3.0 ×*g* 5 min), washed with 1 mL sterile water, and pelleted again. The pellets were resuspended in 410 µL of a transformation mixture containing PEG3350 (30% (w/v) final concentration), lithium acetate (90 mM final concentration), boiled salmon sperm carrier DNA (0.25 mg/mL final concentration), and 2 µg of DNA. For RM-Ec033, 2 µg of the purified plasmid were linearized by digestion for 1 h with *Sca*I, generating blunt ends containing homologous regions to p1. The cells were incubated at 30 °C for 30 min with shaking (200 rpm). Cells were heat shocked for 20 min at 42 °C. The cells were pelleted by centrifugation (3.0 ×*g*, 5 min), resuspended in sterile 0.9% (w/v) NaCl, and plated onto selective SC media minus the appropriate amino acids and NAT selection agar plates. Transformants were grown at 30 °C on plates (3 d for CEN/ARS transformations, 5 d for p1 transformations). Colonies were picked and grown in liquid selective SC medium at 30 °C with shaking (200 rpm) for 2 d until saturation.

**Cytoplasmic plasmid extraction**

Saturated cultures of the yTAH001 transformants grown in selective SC medium were subjected to cytoplasmic plasmid extraction as described in Ravikumar, *et al.*[5] In brief, p1, p2, and p1-derived plasmids were extracted following yeast DNA miniprep. Following p1 transformation, 1.5 mL of saturated culture of cells were pelleted by centrifugation (5.0 ×*g*, 1 min), washed with 1 mL sterile 0.9% (w/v) NaCl, and resuspended in Zymolase solution (US Biological). Cells were incubated at 37 °C for 1 h with rotation (~10 rpm). Cells were pelleted by centrifugation and resuspended in proteinase K (200 µg/mL Sigma) supplemented with 10% SDS and incubated at 60 °C for 30 min with rotation (~10 rpm). Potassium acetate solution (5 M) was added to the solution and placed on ice for 30 min. The solution was centrifuged, and ethanol was added to the supernatant. The solution was pelleted, and the pellet was dried for 30 min. The pellet was resuspended in TE buffer and pelleted by centrifugation. RNase A (1 mg/mL) was added to the supernatant and incubated at 37 °C for 30 min. Isopropanol (1 vol) was added and the solution was pelleted. Integration of p1 into the yeast strain was confirmed via agarose gel electrophoresis of extracted plasmids (Figure S2).

**Plating and screening**

Yeast strain yTAH001 was serially passaged 1:100 in SC-HLUWMC media. After three passages, the culture was plated onto SC-HLUWMC agar plates. After 4 d of colony formation, 94 colonies were picked and inoculated into SC-HLUW media in 96 deep square well plates (500 µL/well). The plate was grown at 30 °C with shaking (200 rpm) overnight. The cells were pelleted by centrifugation (3.0 ×*g*, 5 min) and washed twice with 0.9% NaCl (200 µL). The cells were resuspended in SC-HLUW media containing galactose (20 g/L) in replacement of dextrose. The plate was incubated at room temperature (24 °C) with shaking (200 rpm) for 48 h. The cells were pelleted by centrifugation at 3.0 ×*g* for 5 min and resuspended in PBS (600 µL). Cells were plated in two identical 96-well black plates (Greiner Bio-One, 100 µL/well) and 96-well clear plate (100 µL/well). Coelenterazine diluted in PBS was added to each well (1 µM final concentration). Imaging was performed as described in 'General bioluminescence imaging.' The $OD_{600}$

(absorbance at 600 nm) was measured in a BioTek Synergy H1 microplate reader. Plates were imaged at 1 s exposure for a 5-min sequence. The 8 “hits” with the brightest photon flux fold over the positive control yeast was selected for the next passage. The hits were plated as described previously and the screening cycle was repeated.

**General bioluminescence imaging**

Bioluminescence analyses were performed in solid black, flat-bottom, 96-well plates (Greiner Bio-One). Samples were plated and measured in triplicate unless otherwise stated. Screening plates were imaged in a light-proof chamber using an IVIS Lumina (Xenogen) system with a supercooled CCD camera chilled to -85 °C. The stage was kept at 37 °C during imaging. Living Image software was used to control the instrument and measure photon flux values from defined regions of interest. Exposure time was set to 1 s in all experiments. All other bioluminescence analyses were performed with the Tecan Spark M10 multimode microplate reader. Data was acquired at an integration time of 1000 ms unless otherwise stated. Light emission was measured as counts/s. In yeast experiments, light emission was normalized to $OD_{600}$. For spectral scans, intensities were normalized. All $OD_{600}$ measurements were all acquired on a BioTek Synergy H1 microplate reader (BioTek, Vermont, USA). Fluorescence measurements were acquired on a BioTek Synergy H1 microplate reader (BioTek, Vermont, USA) or a Tecan Spark M10 multimode microplate reader. All data were exported to Microsoft Excel or GraphPad Prism (version 7.0c for Macintosh, GraphPad Software, La Jolla, CA, USA; www.graphpad.com) for further analyses.

**In silico docking**

The effect of the mutations found during evolution on bioluminescence was analyzed by evaluating the catalytic site of NanoLuc. The AutoDock Vina algorithm and the available PDB structure of NanoLuc (PDB: 5IBO) was used to dock CTZ into the active site. Constraints were adjusted as described in Altamash *et al*.[7] A grid box with dimensions of 28 × 28 × 28 with 1 Å spacing and an exhaustiveness of 32 was used. Structural analysis was performed using PyMOL (The PyMOL Molecular Graphics System, Version 2.5.4, Schrödinger, LLC; www.pymol.org; New York, NY, USA).

**Bioluminescence imaging in yeast for variant comparison**

Yeast cultures were grown from glycerol stocks in selective SC media for 2 d at 30 °C. Cultures were pelleted (3.0 ×*g*, 5 min) and washed with 0.9% NaCl twice. The pellets were resuspended in PBS and all cultures were diluted to the same $OD_{600}$ (~0.2). Cultures were plated in triplicate (50 µL/well) in black 96-well plates. Fluorescence of mNeonGreen (excitation 485 nm, emission 517 nm, 15 nm bandwidth) was measured on a Tecan Spark M10 multimode microplate reader. Luminescence was then acquired on Tecan F200 Pro injection port luminometer with a neutral density filter. CTZ was prepared as a 2X stock solution in PBS, and 100 µL was added to assay wells. Plates were imaged 2.5 min before luciferin addition, and continuously every 30 s for 15 min after injection. Luminescence emission was normalized to mNeonGreen expression.

**Protein expression and purification**

Proteins were encoded in a pCold vector. Glycerol stocks for *E. coli* BL21 cells expressing the proteins of interest were streaked onto LB plates containing ampicillin (100 µg/mL). Single colonies were picked into starter cultures of LB Broth (5 mL) containing ampicillin (100 µg/mL) and grown overnight (37 °C, 220 rpm). Starter culture (250 µL) was used to inoculate 10 mL of LB Broth. Cultures were grown at 37 °C at 220 rpm until reaching an optical density ($OD_{600}$) of ~0.8 and cold-shocked at 4 °C for 1 h. The cultures were induced with 0.5 mM isopropyl-β-D-thiogalactopyranoside (IPTG) and shaken (220 rpm) at 16 °C for 18 h. The cultures were harvested (4000 rpm, 10 min) and resuspended in lysis buffer (50 mM Tris HCl, 150 mM NaCl, 0.5% Tween-20, pH 7.4). Protease inhibitor phenylmethylsulfonyl fluoride (PMSF, 0.1 mM) was added to the cell resuspension. The resuspension was sonicated (QSonica) at 40% amplitude, using 2 sec on/2 sec off intervals for 15 min. The cultures were centrifuged to separate cell debris (10,000 rpm, 10 min). The clarified lysate supernatant was purified by Ni-NTA chromatography. Columns were washed with wash buffer (20 mM imidazole, 50 mM sodium phosphate, pH 7.4). Proteins were eluted with elution buffer (200 mM imidazole, 50 mM, sodium phosphate, pH 7.4). Eluted proteins were concentrated via spin column (GE Healthcare, Vivaspin, 5 kDa MWCO) and buffer exchanged into sodium phosphate buffer (50 mM, pH 7.4). SDS-PAGE analyses were performed and gels were stained with Coomassie Brilliant Blue R-250. Final protein concentrations were determined by BCA assay. Proteins were stored in 50% glycerol at –20 °C. For all bioluminescence imaging, the purified recombinant protein was diluted in PBS.

**Kinetic analysis**

Measurements were acquired on a Tecan F200 Pro injection port luminometer with a neutral density filter. Reactions were performed in black 96-well flat-bottom plates (Grenier). Solutions of CTZ (0.05-10 µM, in PBS) were prepared and added (100 µL) to each well. The resulting luminescence was measured for 1.5 s prior to the addition of enzyme (0.1 nM, 100 µL) in PBS. Following the addition of enzyme, luminescence was recorded every 0.2 s over a 60 s period. Samples were analyzed in triplicate. The peak intensities were determined by averaging the three maximum photon outputs per run. Km and relative kcat values were determined using nonlinear regression analyses in Prism (GraphPad).

**General cell culture methods**

Human embryonic kidney (HEK293T) cells (ATCC, Manassas, VA, USA) were cultured in complete media: Dulbecco's Modified Eagle's Medium (DMEM, Corning) containing 10% (v/v) fetal bovine serum (FBS, Life Technologies), penicillin (100 U/mL) and streptomycin (100 µg/mL, Gibco), and Plasmocin® prophylactic (5 µg/mL). Cells were incubated at 37 °C in a 5% $CO_2$ humidified chamber. Cells were serially passaged using trypsin (0.25 % in HBSS, Gibco).

**In cellulo bioluminescence imaging**

HEK293T cells were plated in a 12-well plate in complete media (1 mL, 100,000 cells/well). Following incubation at 37 °C overnight, the cells were transiently transfected with 1000 ng of DNA (Addgene #85200 or pTAH061) using cationic lipid formulations (Lipofectamine 3000; Invitrogen). After 24 h, the cells were lifted with trypsin, neutralized with media, and pelleted by centrifugation (500 *g*, 5 min). The cells were washed twice with PBS. The cell pellet was resuspended in PBS, and 10,000 cells/well were plated in triplicate (50 µL/well) in black 96-well plates. Measurements were acquired on a Tecan F200 Pro injection port luminometer with a neutral density filter. CTZ was prepared as a 2X stock solution in PBS, and 100 µL was added to assay wells. Plates were imaged 2.5 min before luciferin addition, and continuously every 30 s for 20 min after injection. Mean fluorescence intensity (MFI) was analyzed via mNeonGreen expression and measured 1 d post-transfection via flow cytometry.

**Flow cytometry**

Cells were trypsinized and washed twice in PBS prior to analysis on a ACEA NovoCyte Flow Cytometer. For each sample, 100,000 live cell events were collected. Data were analyzed using NovoExpress software (version 1.3.0) and FlowJo (version 10.10.0).

**Supplementary Table S1. Mutated residues found in GeNL from top hits in evolution.**

| | base pairs 1057-1059 | | base pairs 1348-1350 | | base pairs 1555-1557 | |
|---|---|---|---|---|---|---|
| **sequence** | **codon** | **amino acid** | **codon** | **amino acid** | **codon** | **amino acid** |
| GeNL sequence | GAT | D | TTA | L | CAT | H |
| passage 0 | GAT | D | TTA | L | CAT | H |
| passage 1, mutant 73 | TAT | Y | ATA | I | TAT | Y |
| passage 2, mutant 84 | TAT | Y | ATA | I | TAT | Y |
| passage 3, mutant 47 | TAT | Y | ATA | I | TAT | Y |
| passage 4, mutant 58 | TAT | Y | ATA | I | TAT | Y |
| passage 5, mutant 68 | TAT | Y | ATA | I | TAT | Y |
| passage 6, mutant 60 | TAT | Y | ATA | I | TAT | Y |
| passage 7, mutant 36 | TAT | Y | ATA | I | TAT | Y |

The top mutant from each passage was sequenced by bulk Sanger sequencing and compared to the GeNL sequence in passage 0, the starting point for evolution. Only the base pair mutations resulting in an amino acid change in GeNL are shown. Mutated base pairs are shown in red.

**Supplementary Table S2. Enzymatic characteristics of luciferase mutants with CTZ.**

| Luciferase | $K_m$ (μM) | $k_{cat}/K_m$ | $R^2$ |
|---|---|---|---|
| GeNL | 1.93 ± 0.17 | $2.81 \times 10^9$ | 0.99 |
| L243I | 2.98 ± 0.70 | $1.69 \times 10^9$ | 0.92 |
| L243I/H312Y | 4.12 ± 0.32 | $7.64 \times 10^9$ | 0.99 |

**Supplementary Table S3. Key plasmids used in this study.**

| Plasmid name | Description | Expression in | Reference |
|---|---|---|---|
| pCEN/ARS-pREV1-BadBoy2 | A CEN/ARS plasmid encoding the BadBoy2 error-prone DNAP sequence under the pREV1 promoter. | *S. cerevisiae* | Rix *et al.*[8] |
| pGKL1-short-M17 | Landing pad for p1: p1:ShortPol::p2O5-MET17 | *S. cerevisiae* | Wellner *et al.*[3] |
| RM-Ec033 | p10B2-GeNL yeast codon optimized-Aga2p for generation of an integration cassette onto p1 for surface display | *S. cerevisiae* | This work |
| pTAH003 | pCold-GeNL | *E. coli* | This work |
| pTAH004 | pCold-L243I | *E. coli* | This work |
| pTAH005 | pCold-H312Y | *E. coli* | This work |
| pTAH006 | pCold-GeNL | *E. coli* | This work |
| pTAH023 | CEN/ARS-GeNL | *S. cerevisiae* | This work |
| pTAH024 | CEN/ARS-L243I | *S. cerevisiae* | This work |
| pTAH025 | CEN/ARS-H312Y | *S. cerevisiae* | This work |
| pTAH026 | CEN/ARS-L243I/H312Y | *S. cerevisiae* | This work |
| Addgene #85200 | pcDNA-GeNL | mammalian | Suzuki *et al.*[9] |
| pTAH061 | pcDNA-GeNL-L243I | mammalian | This work |

**Supplementary Table S4. Key yeast strains used in this study.**

| Yeast strain | Description | Genome |
|---|---|---|
| yAW723 | galactose inducible AHEAD strain with BadBoy2 polymerase encoded on a CEN/ARS nuclear plasmid | MATa AGA1::GAL1AGA1::URA3 ura352 trp1 leu2Δ200 his3Δ200 pep4::HIS3 prb11.6R can1 GAL Trp1:: Δ-0 Met17:: Δ-0 p1:ShortPol::p2O5-MET17 CEN/ARS-pREV1-BadBoy2 |
| yTAH001 | galactose inducible AHEAD strain with BadBoy2 polymerase encoded on a CEN/ARS nuclear plasmid and GeNL integrated onto p1 for surface display | MATa AGA1::GAL1AGA1::URA3 ura352 trp1 leu2Δ200 his3Δ00 pep4::HIS3 prb11.6R can1 GAL Trp1:: Δ-0 Met17:: Δ-0 p1:ShortPol::p2O5-MET17 CEN/ARS-pREV1-BadBoy2; p1: RM-Ec033 |
| yAW680 | galactose inducible yeast surface display strain with OrthoRep landing pad | EBY100 Trp1D0 met17D0 p1-shortpol-MET17 |
| yTAH005 | Yeast strain with GeNL encoded on a CEN/ARS nuclear plasmid | EBY100 Trp1D0 met17D0 p1-shortpol-MET17; CENARS: pTAH023 |
| yTAH006 | Yeast strain with GeNL-L243I encoded on a CEN/ARS nuclear plasmid | EBY100 Trp1D0 met17D0 p1-shortpol-MET17; CENARS: pTAH024 |
| yTAH007 | Yeast strain with GeNL-H312Y encoded on a CEN/ARS nuclear plasmid | EBY100 Trp1D0 met17D0 p1-shortpol-MET17; CENARS: pTAH025 |
| yTAH008 | Yeast strain with GeNL-L243I/H312Y encoded on a CEN/ARS nuclear plasmid | EBY100 Trp1D0 met17D0 p1-shortpol-MET17; CENARS: pTAH026 |

**Supplementary Table S5. Primers constructed for this study.**
Primers are listed 5′ → 3′.

| Primer name | Purpose | Primer sequence |
|---|---|---|
| TAH03_Fwd | forward primer for amplification of gene insert for p1 | ATAGGGGAGAGTACTAAAAGTGAG |
| TAH05_Fwd | GeNL sequencing primer | CGCTTCTATCGCTGCTAAGGAAG |
| TAH06_Fwd | Gibson overhang forward primer for GeNL [Nagai] in pCold vector | GCATCATCATCATCATCATATCGAAGGTAGGCATATGATGGTGTCCAAGGGCGAAGAG |
| TAH07_Rev | Gibson overhang reverse primer for GeNL [Nagai] in pCold vector | CTATCTAGACTGCAGGTCGACAAGCTTGAATTCTTACGCCAGAATGCGTTCGCACAG |
| TAH08_Rev | reverse primer for amplification of gene insert for p1 | CTTCTTCAAAAAACATACTGTGTGTTTATG |
| TAH09_Rev | reverse primer for SDM (L243I) on GeNL | GACTTGGTCAATGTTGTAGCCGGCTGTCTGTCGCCAGTCCCCAACGAAATC |
| TAH10_Fwd | forward primer for SDM (L243I) on GeNL | GGCTACAACATTGACCAAGTCCTTGAACAGGGAGGTGTGTCCAGTTTG |
| TAH11_Rev | reverse primer for SDM (H312Y) on GeNL | CACCTTAAAATAATGATCATCCACAGGGTACACCACCTTAAAAATTTTTTCGATC |
| TAH12_Fwd | forward primer for SDM (H312Y) on GeNL | GGATGATCATTATTTTAAGGTGATCCTGCACTATGGCACACTGGTAATCGAC |
| TAH013_Fwd | forward GeNL amplicon for Golden Gate (base pairs 1-747) | GCATCGTCTCATCGGTCTCATATGGTGTCAAAAGGCGAAGAAGACAAC |
| TAH014_Rev | reverse GeNL amplicon for Golden Gate (base pairs 1-747) | CGTCTCAAATATTGTACCCCGCAGTCTGAC |
| TAH015_Fwd | forward GeNL amplicon for Golden Gate (base pairs 1-747) | CGTCTCATATTGATCAAGTCTTGGAACAAGGAGGT |
| TAH016_Rev | reverse GeNL amplicon for Golden Gate (base pairs 1-747) | CGTCTCAATAATGATCGTCGACTGGGTAAACAAC |
| TAH017_Fwd | forward GeNL amplicon for Golden Gate (base pairs 1-747) | CGTCTCATTATTTCAAAGTTATCCTACACTATGGCACC |
| TAH018_Rev | reverse GeNL amplicon for Golden Gate (base pairs 1-747) | ATGCCGTCTCAGGTCTCAGGATAGCCAATATTCGCTCGCATAGA |
| TAH029_Fwd | Gibson overhang forward primer for GeNL in CEN/ARS vector | CATATCCAGCGAGATCTATGGTGTCAAAAGGCG |
| TAH030_Rev | Gibson overhang reverse primer for GeNL in CEN/ARS vector | GAAATTCGCCTCGAGTTAAGCCAATATTCGCTCG |
| ZY28 | Gibson overhang forward primer for GeNL in pcDNA vector | ATACGACTCACTATAGGGAGACCCAAGCTTCGCCACCATGGTGTCCAAGGGCGAAGAGGA |
| ZY29 | Gibson overhang reverse primer for GeNL in pcDNA vector | GCCGCCAGTGTGATGGATATCTGCAGAATTCCTACGCCAGAATGCGTTCGC |

**Supplementary Note S1.** Plasmid sequence for RM-Ec033.

Full sequence for the plasmid used to integrate onto p1:
GeNL
Aga2p

gagatacctacagcgtgagctatgagaaagcgccacgcttcccgaagggagaaaggcggacaggtatccggtaagcggcaggg
tcggaacaggagagcgcacgagggagcttccagggggaaacgcctggtatctttatagtcctgtcgggtttcgccacctctgacttga
gcgtcgatttttgtgatgctcgtcaggggggcggagcctatggaaaaacgccagcaacgcggcctttttacggttcctggccttttgctgg
ccttttgctcacatgttctttcctgcgttatcccctgattctgtggataaccgtattaccgcctttgagtgagctgataccgctcgccgcagccg
aacgaccgagcgcagcgagtcagtgagcgaggaagcggaagagcgcccaatacgcaaaccgcctctccccgcgcgttggccg
attcattaatgcagctggcacgacaggtttcccgactggaaagcgggcagtgagcgcaacgcaatagtactataatatatgaattaca
ttattaatttaaaaacgcgtgatgacctatacataggaagatctatagaaacaaaaagattaataactttcaaatatcagaaaaatatag
aaacatgtgataagctcatagacatAtaaaggatccATGAGATTCCCATCTATCTTCACCGCTGTTTTGTTC
GCTGCTTCTTCTGCTTTGGCTGCTCCAGCTAACACCACCACCGAAGACGAAACCGCTCAAA
TCCCAGCTGAAGCTGTTATCGACTACTCTGACTTGGAAGGTGACTTCGACGCTGCTGCTTT
GCCATTGTCTAACTCTACCAACAACGGTTTGTCTTCTACCAACACCACCATCGCTTCTATCG
CTGCTAAGGAAGAAGGTGTTCAATTGGACAAGAGAGAAGCTGACGCAggtaccGTGTCAAAA
GGCGAAGAAGACAACATGGCATCGCTACCCGCAACGCATGAACTACATATTTTCGGTTCAA
TAAACGGTGTGGATTTTGACATGGTTGGTCAGGGTACTGGAAACCCCAACGACGGATATGA
AGAACTAAACTTAAAGTCCACAAAGGGTGACCTACAATTTTCACCGTGGATCCTGGTCCCA
CATATTGGTTATGGGTTTCATCAGTATCTCCCATATCCGGATGGTATGTCCCCATTTCAAGC
CGCTATGGTAGATGGATCCGGGTATCAAGTACATAGGACAATGCAATTCGAAGATGGAGCT
TCTTTAACTGTCAACTACAGATACACCTACGAAGGTAGTCATATTAAAGGAGAAGCTCAAGT
AAAAGGGACAGGTTTCCCAGCGGATGGCCCAGTTATGACAAACAGTTTAACAGCTGCCGA
TTGGTGTAGATCCAAGAAGACATACCCCAATGATAAGACTATCATTAGTACCTTCAAATGGA
GCTACACAACCGGTAATGGTAAGAGATATAGATCTACTGCAAGAACCACGTATACGTTTGC
TAAACCAATGGCGGCTAACTATCTTAAAAACCAACCGATGTATGTCTTTCGAAAGACTGAAT
TAAAGCATTCTAAGACAGAATTGAACTTCAAAGAATGGCAGAAGGCTTTCACGGGTTTTGAA
GATTTTGTAGGTGATTGGCGTCAGACTGCGGGGTACAATTTAGATCAAGTCTTGGAACAAG
GAGGTGTGTCTTCGTTGTTTCAAAACTTGGGTGTGTCCGTCACCCCAATACAAAGAATAGT
TTTATCCGGGGAAAACGGTCTGAAGATCGATATTCATGTCATTATCCCATACGAGGGCCTT
AGTGGGGATCAGATGGGTCAAATTGAGAAAATTTTTAAGGTTGTTTACCCAGTCGACGATC
ATCATTTCAAAGTTATCCTACACTATGGCACCCTTGTTATTGATGGTGTTACTCCAAACATG
ATTGATTATTTCGGGAGGCCCTACGAAGGAATCGCAGTTTTCGATGGGAAAAAAATTACGG
TTACAGGTACTCTATGGAACGGGAATAAGATTATTGACGAAAGATTAATCAATCCAGATGGT
TCATTGTTGTTTAGAGTAACTATCAATGGTGTGACTGGTTGGCGTCTATGCGAGCGAATATT
GGCTgtcgacGGTTCCaaggacaatagctcgacgattgaaggtagatacccatacgacgttccagactacgctctgcaggct
agtggtggtggtggttctggtggtggtggttctggtggtggtggttctgctagcgacgtcCAGGAACTGACAACTATATGCG
AGCAAATCCCCTCACCAACTTTAGAATCGACGCCGTACTCTTTGTCAACGACTACTATTTTG
GCCAACGGGAAGGCAATGCAAGGAGTTTTTGAATATTACAAATCAGTAACGTTTGTCAGTA
ATTGCGGTTCTCACCCCTCAACAACTAGCAAAGGCAGCCCCATAAACACACAGTATGTTTT
TTGAAGAAGACTTGtcatgaAAAAAAAAAAAAAAAAAAAAAAAAAAAAAAAAAAAAAAAAAAAAA
AAAAAAAAAAAAAAAAAAAAAAAAAAAAAACCTGTCACCGGATGTGTTTTCCGGTCTGATGA
GTCCGTGAGGACGAAACAGGgaattcgcgaatttcttatgatttatgatttttattattaaataagttataaaaaaaataagtgt
atacaaattttaaagtgactcttaggttttaaaacgaaaattcttattcttgagtaactctttcctgtaggtcaggttgctttctcaggtatagcat
gaggtcgctcgagctcttaagggcaaggcatagacatatacaaggcctgttcaccgtcagatgctgttccatcatataaagcagtatct
aaaccacataatgtgaaacccattcttctataagcatggatagcaggggcgttaacattagtaacttctaaccataaatgaccggctcct
ctctctcttgcaaattctgtagccaaacccatcaaagctcttcctacaccatgacctctatgctctggggcgacttctatatcttcaacggtc
aaccttctattccaacctgaataagaaacgactacaaaaccagccaaatcaccgtcatcaccgtatgctacgaaagttcttgaatctgg
gtcaccatcttctccgtcgtcgctttcgtcatcagattcatcatctggaaagactttagttaatggaggatcgactgggacttctcttaaggta

aatccatcaccggtggcggttactctaaagacggtgtcggtggtgaatgaaccgtctaaagcctcaattgcttcagcatcaccaggga
ctgatgttctgtatctgtaggctgtatcatctaaggtagtacccattttataattattataaaaactttcatatagagctccgtttctattatgaattt
catttataaagtttatgtacaaatatcataaaaaaagagaatctttttttcttagcatttttgacgaaatttgctattttgttagagtcttttacacc
atttgtctccacacctccgcttacatcaacaccaataacgccatttaatctaagcgcatcaccaacattttctggcgtcagtccaccagct
aacataaaatgtaagctttcggggctctcttgccttccaacccagtcagaaatcgagttccaatccaaaagttcacctgtcccacctgctt
ctgaatcaaacaagggaataaacgaatgaggtttctgtgaagctgcactgagtagtatgttgcagtcttttggaaatacgagtcttttaat
aactggcaaaccgaggaactcttggtattcttgccacgactcatctccatgcagttggacgatatcaatgccgtaatcattgaccagag
ccaaaacatcctccttaggttgattacgaaacacgccaaccaagtatttcggagtgcctgaactatttttatatgcttttacaagacttgaa
attttccttgcaataaccgggtcaattgttctctttctattgggcacacatataatacccagcaagtcagcatcggaatctagagcacattct
gcggcctctgtgctctgcaagccgcaaactttcaccaatggaccagaactacctgtgaaattaataacagacattttataattattataaa
aactttcatataagagatataaaatttaatatggaaaaaataagacaagaaagatacaaccaaatgaaagaagctctaaatagtgtt
gaaggttataaaggaaaaattgtagcctcagactcagattggtgtttcaaagatcctCAAGGCAATAGAATAACAGATTT
TGATAGTactttaagccagccccgacacccgccaacacccgctgacgcgccctgacgggcttgtctgctcccggcatccgcttac
agacaagctgtgaccgtctccgggagctgcatgtgtcagaggttttcaccgtcatcaccgaaacgcgcgagacgaaaggcaaaaa
aaaaggctccaaaaggagcctttaattgtatcggtttaccaatgcttaatcagtgaagcccctatttcagcaatctgtctatttcgttcgtcc
atagttgcctgactacccgtcgtgtagataactacgatacgggaaggcttaccatctggtccaagtgcggcaatgataccccgtgaac
cacgctctcctgccccagatttatcagcaataaaccagccagcaggaagtgccgaacgcagtagtggtcctgcaactttatccgcttc
aagccagtctattaattgttgccgtgaagccagagtaagtagttcgccagttaatagtttgcgtaatgttgttgccattgccgcaggcattgt
ggtgtctcgctcgtcgtttggtatggcttcattcagttctggttcccaacggtcaaggcgagttacatggtctcccatattgtgcaaaaaagc
ggttaattccttcggtccaccgatggttgtcagaagtaagttcgccgcagtgttatcactcatcgttatggcggcactacataattctcttac
cgtcattccgtccgtaaggtgcttttctgtcactggggagtattcaaccaagtcattctgcgaatagtgtattcgtcgtccgagttgctcttgc
cccgcatctacacgggataatactgctccacatagcagaactttgaaagtgctcatcattggaaaacgctcttcagggcgaaaactctc
aaggattttaccgctattcaagtccagttctatgtatcccactcgtgcccccaattggtcctctgcgtcttttactttcacgagcgtttctgggtg
agcaaaaacaggaaggcaaaatgccgcaaaaaagggaataagggcgacacggaaatgttgaatactcatactcttcctttttcaat
attattgaagcatttatcagggttattgtctgatgagcggatacatatttgaatgtatttagaaaaataaacaaataggggaattaaaaaa
aagcccgctcattaggcgggctactagtcggatatagttcctcctttcagcaaaaaacccctcaagacccgtttagaggccccaaggg
gttatgctagttattgctcagcggtggcagcagccaactcagcttcctttcgggctttgttagcagccggatcttctagaagaccccgtag
aaaagatcaaaggatcttcttgagatcctttttttctgcgcgtaatctgctgcttgcaaacaaaaaaaccaccgctaccagcggtggtttgt
ttgccggatcaagagctaccaactctttttccgaaggtaactggcttcagcagagcgcagataccaaatactgttcttctagtgtagccgt
agttaggccaccacttcaagaactctgtagcaccgcctacatacctcgctctgctaatcctgttaccagtggctgctgccagtggcgata
agtcgtgtcttaccgggttggactcaagacgatagttaccggataaggcgcagcggtcgggctgaacggggggttcgtgcacacagc
ccagcttggagcgaacgacctacaccgaact
